# Identifying differential isoform abundance with RATs: a universal tool and a warning

**DOI:** 10.1101/132761

**Authors:** Kimon Froussios, Kira Mourão, Gordon G. Simpson, Geoffrey J. Barton, Nick J. Schurch

**Affiliations:** Division of Computational Biology, School of Life Sciences, University of Dundee, Dundee, UK; Centre for Gene Regulation & Expression, School of Life Sciences, University of Dundee, Dundee, UK; Division of Plant Sciences, School of Life Sciences, University of Dundee, Dundee, UK; Cell and Molecular Sciences, James Hutton Institute, Invergowrie, Dundee, UK

## Abstract

**Motivation:** The biological importance of changes in gene and transcript expression is well recognised and is reflected by the wide variety of tools available to characterise these changes. Regulation via Differential Transcript Usage (DTU) is emerging as an important phenomenon. Several tools exist for the detection of DTU from read alignment or assembly data, but options for detection of DTU from alignment-free quantifications are limited.

**Results:** We present an R package named *RATs* – (Relative Abundance of Transcripts) – that identifies DTU transcriptome-wide directly from transcript abundance estimations. *RATs* is agnostic to quantification methods and exploits bootstrapped quantifications, if available, to inform the significance of detected DTU events. *RATs* contextualises the DTU results and shows good False Discovery performance (median FDR ≤0.05) at all replication levels. We applied *RATs* to a human RNA-seq dataset associated with idiopathic pulmonary fibrosis with three DTU events validated by qRT-PCR. *RATs* found all three genes exhibited statistically significant changes in isoform proportions based on Ensembl v60 annotations, but the DTU for two were not reliably reproduced across bootstrapped quantifications. *RATs* also identified 500 novel DTU events that are enriched for eleven GO terms related to regulation of the response to stimulus, regulation of immune system processes, and symbiosis/parasitism. Repeating this analysis with the Ensembl v87 annotation showed the isoform abundance profiles of two of the three validated DTU genes changed radically. *RATs* identified 414 novel DTU events that are enriched for five GO terms, none of which are in common with those previously identified. Only 141 of the DTU evens are common between the two analyses, and only 8 are among the 248 reported by the original study. Furthermore, the original qRT-PCR probes no longer match uniquely to their original transcripts, calling into question the interpretation of these data. We suggest parallel full-length isoform sequencing, annotation pre-filtering and sequencing of the transcripts captured by qRT-PCR primers as possible ways to improve the validation of RNA-seq results in future experiments.

**Availability:** The package is available through Github at https://github.com/bartongroup/Rats.

## Introduction

High-throughput transcriptomics experiments have until recently focused on quantifying gene expression and calculating Differential Gene Expression (DGE) between samples in different groups, conditions, treatments, or time-points. In higher eukaryotes, alternative splicing of multi-exon genes and alternative transcript start and end sites can lead to multiple transcript isoforms originating from each gene. Since isoforms can have different and distinct functions [1–3], analysis of Differential Transcript Expression (DTE) is preferable to DGE. Unfortunately, isoform-level transcriptome analysis is more complex and expensive since, in order to achieve similar statistical power in a DTE study, higher sequencing depth is required to compensate for the expression of each gene being split among its component isoforms. In addition, isoforms of a gene typically have a high degree of sequence similarity and this complicates the attribution of reads among them. Despite these challenges, several studies have shown that shifts in individual isoform expression represent a real level of gene regulation with phenotypic consequences [4–7], suggesting there is little justification for choosing DGE over DTE in the study of complex transcriptomes.

It is possible to find significant DTE among the isoforms of a gene, even when the gene shows no significant DGE. This introduces the concept of Differential Transcript Usage (DTU), where the individual isoform abundances of a gene can change relative to one another, with the most pronounced examples resulting in a change of the dominant isoform (isoform switching). The definitions of DGE, DTE and DTU are illustrated in Figure 1.

**Figure 1.**
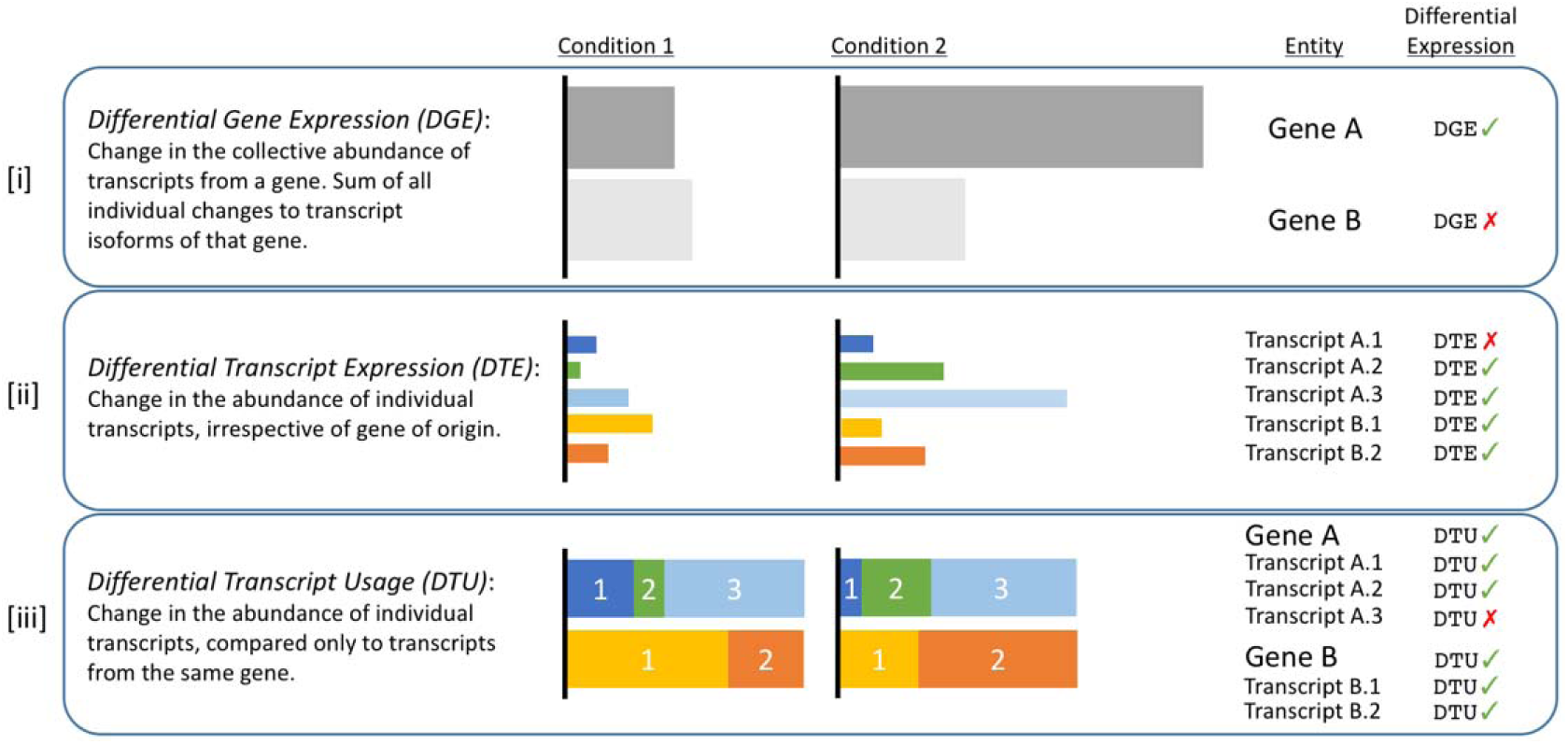
*Illustrative definitions of the three types of differential expression analysis (DGE, DTE and DTU) for two genes (Gene A and Gene B), with 3 and 2 isoforms respectively, whose expression is compared across two conditions (Condition 1 and Condition 2). The horizontal width of each coloured box represents the abundance of the relevant gene or transcript. A negative differential expression result (red cross-mark) for a given entity in any one of the three analysis types does not exclude that same entity from having a positive result (green tick-mark) in one of the other two analysis types. [iii] The relative isoform abundances are scaled to the absolute isoform abundances [ii], which in turn are scaled to the gene expressions in [i]. Gene A is differentially expressed, but only two of its three isoforms are differentially expressed (A.2 and A.3). Proportionally, Gene A’s primary isoform (A.3) remains the same, but the ratios of the two less abundant isoforms change. Gene B is not differentially expressed, but both its isoforms are differentially expressed, and demonstrate an example of isoform switching.*

To quantify the isoforms and assess changes in their abundance, most existing tools for DTU analysis (e.g. Cufflinks [8], DEXSeq [9], LeafCuttter [10]) rely on reads that either span splice-junctions or align to unique exons. However, with the newest generation of transcript quantification tools (Kallisto [11], Sailfish [12], Salmon [13]), reads are not aligned to the transcriptome or the genome. Instead, these tools combine a pseudo-mapping of the *k*-mers present within each read to the *k*-mer distributions from the transcript annotation with an expectation maximization algorithm, to infer the expression of each transcript model directly. Alignment-free methods are much faster than traditional alignment-based methods (RSEM [14], TopHat2 [15], STAR [16]) or assembly-based methods (Cufflinks [8], Trinity [17]), but the lack of alignments prevents these new methods from being compatible with differential expression methods such as Cufflinks, DEXSeq and Leafcutter. Instead, Sleuth [18] is a tool that handles DTE analysis from alignment-free transcript quantifications, but DTU analysis is currently less straight-forward. SwitchSeq [19] and iso-kTSP [6] focus on a particular subset of DTU analysis from alignment-free data, namely isoform switching. SUPPA [20,21], on the other hand, identifies differential splicing events.

In this paper, we present *RATs*, a tool for identifying DTU directly from isoform quantifications. It is designed primarily for use with alignment-free abundance data and can take advantage of bootstrapped quantifications, but remains agnostic to the quantification method and can also be used in alignment-based workflows. We compare the results from *RATs* to published and validated instances of DTU and demonstrate that the results of both RNA-seq based and qRT-PCR based analyses are sensitive to the annotation used for transcript quantification and primer design, respectively.

## Implementation

*RATs* applies filters to the data prior to any statistical testing in order to reduce both the number of low quality calls and the number of tests carried out. These filters are (i) isoform ratio changes can only be defined for genes that are expressed in both conditions and (ii), transcript abundances need to exceed a minimum count threshold. The workflow of *RATs* is illustrated in Figure 2.

**Figure 2.**
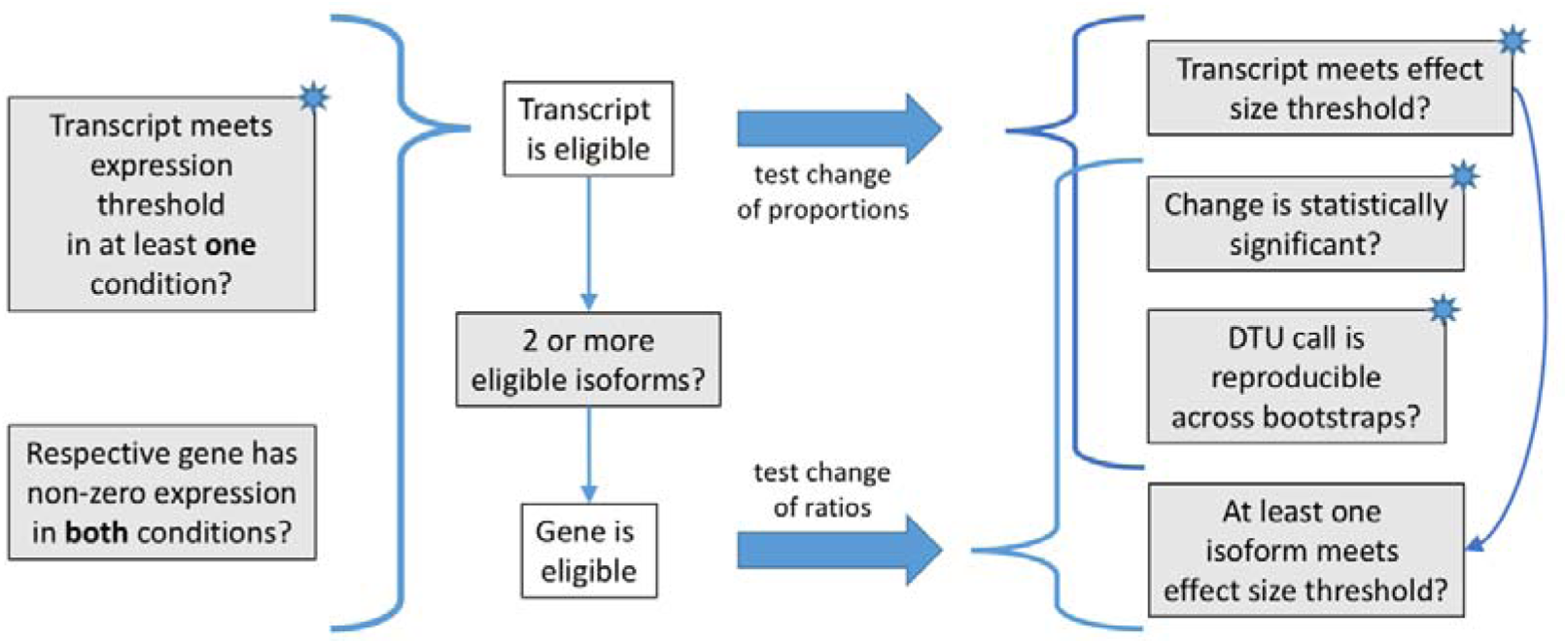
*RATs workflow. Individual transcripts whose estimated abundance falls below a specified threshold in both conditions are excluded. Genes that are completely switched off in one or both conditions are also excluded, as isoform proportions cannot be defined for them. The remaining transcripts are tested for significant changes in their individual proportions. Genes with at least two isoforms above the expression threshold are tested for significant change in their expressed isoform ratios. Statistically significant results are then required to exceed a minimum change in isoform proportion and to be reproducible in random bootstrapping of the quantification data. All criteria marked with a star in their upper right corner represent user-defined runtime parameters.*

Significant relative transcript abundance changes are detected using two separate approaches: one detects DTU at the gene level and the other detects DTU at the transcript level. The G-test of independence [22], without continuity corrections, is implemented in *RATs* and used by both approaches. At the gene level, *RATs* compares the set of each gene’s isoform abundances between two conditions to identify if the abundance ratios have changed. At the transcript level, *RATs* compares the abundance of each individual transcript against the pooled abundance of its sibling isoforms to identify changes in the proportion of the gene’s expression attributable to that specific transcript. Both methods include the Benjamini-Hochberg false discovery rate correction for multiple testing [23], and achieve median false positive rates ≤0.05 (*FPR = false positives / input negatives*) even with only three replicates per condition, with notable improvements at higher replication levels, as shown in Figure 3 (panel A).

**Figure 3.**
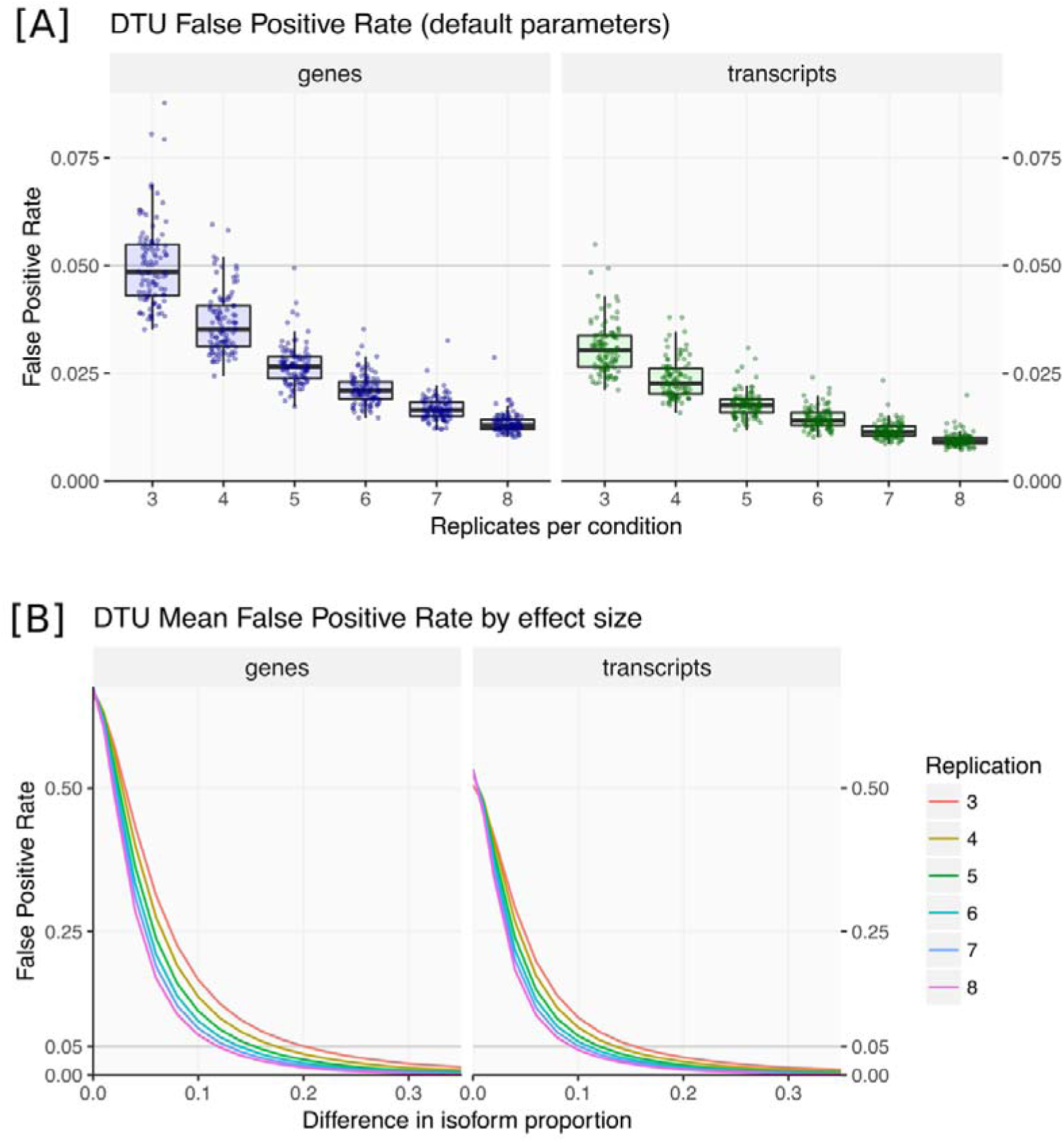
*False positive rate (FPR = False Positives / Input Negatives) of RATs as a function of replication level. FPR measured over 100 bootstrap iterations of randomly selected (without replacement) replicates from a pool of 16 high-quality wild-type Colombia-0 Arabidopsis thaliana replicates from Froussios et al.* [24]*. [A] FPR of each bootstrap iteration, for default values of all RATs parameters (v0.4.4), across a range of replication levels. [B] Mean FPR by replication level, as a function of effect size threshold (effect size = difference between an isoform’s proportions). For genes, the effect size is defined as the largest proportion difference observed among each gene’s isoforms. In every iteration, the FPR was calculated against the number of genes or transcripts that were eligible for testing each time, but that number remained very stable across bootstraps and replication levels (Supplementary file 1). In this case the FPR calculated equates to the False Discovery Rate, because the p-values are appropriately corrected for multiple testing.*

These two approaches to identifying DTU have different strengths and weaknesses when applied to an RNA-seq dataset. Simultaneously utilizing the expression information across all the isoforms in a gene makes the gene-level test sensitive to smaller changes in relative expression, compared to testing transcripts individually, but also more prone to false positives. Figure 3, shows that the gene-level test has a higher mean and median FPR than the transcript-level test irrespective of replication or effect size, although the two methods converge for highly replicated experiments and/or large effect sizes. Furthermore, it is only applicable to genes with at least two isoforms that both pass the pre-filtering criteria imposed by *RATs*, potentially limiting its utility for genes with low read coverage. Finally, the gene-level test only identifies the presence of a shift in the ratios of the isoforms belonging to the gene, without identifying which specific isoforms are affected. The transcript-level test, by contrast, directly identifies the specific isoforms whose proportions are changing and has a lower False Positive Rate than the gene-level test. However, considering each isoform independently requires a larger number of tests to be performed, thus resulting in a greater multiple testing penalty and a lower statistical power. *RATs* currently does not combine the results of the two methods into a single verdict, but rather presents both as complementary views of the data.

Alignment-free transcript quantification is hundreds of times faster than traditional read alignment methods [11,13], making it feasible to repeat the process many times on iterative subsets of the read data. This bootstrapping approach quantifies the technical variance in the transcript abundance estimates. *RATs* provides the option to use these bootstrapped abundance estimates to apply a reproducibility constraint on the DTU calls, by replacing the mean abundances with the values from random individual iterations and measuring the fraction of these iterations that result in a positive DTU classification. This bootstrapping process reveals the extent to which the variability in the quantification of the expression of each transcript affects the DTU identification of the gene and a threshold can be set on the reproducibility of this classifications. Similarly, *RATs* also optionally measures the reproducibility of the DTU results relative to the inter-replicate variation by sub-setting the samples. Finally, two further user-defined thresholds are applied: one filters out low-expression transcripts, while the other excludes transcripts according to the change in their DTU proportion (effect size).

*RATs* is implemented in R [25]. It is distributed through Github as an R source package (https://github.com/bartongroup/RATS) and has been freely available since August 2016. A detailed manual is included in the package.

## Input and output

*RATs* accepts as input either a *Sleuth* object [18] or, alternatively, a set of tables of fragment count estimates (with or without bootstrap information). An annotation mapping the correspondence between transcript and gene identifiers is also required. This can be given directly as a table or inferred from a GTF file.

*RATs* results are returned in the form of R *data.table* objects [26]. Along with the DTU calls, the results record the full provenance of the calculations, allowing the user to trace back the decision steps behind each call and gain better understanding of their data. Additionally, *RATs* employs the *ggplot2* package [27] to provide visualisations of the results. In addition to a “volcano plot” view of the DTU effect size against the significance, *RATs* also visualises the DTU classifications for a given gene in the context of the respective isoform abundances and their consistency across the replicates. An example, drawn from the case study in the following section, is shown in Figure 4. Panel A shows the absolute (upper panels) and relative (lower panels) abundances for each isoform within each condition. Panel B shows the same information, now reorganised to facilitate the comparison of transcript abundance between conditions. From these plots it is straightforward to identify the dominant isoform in each condition and the cases of isoform up-or down-regulation. In this example ENST00000457634 is the dominant transcript from this gene in the control condition but is down-regulated in the IPF condition, while ENST00000490573 is strongly up-regulated in the IPF condition becoming the dominant isoform. The lines connecting transcript abundance points in panel A show the transcript expression profile for this gene for each of the three replicates, making it easy to identify outlier behaviour that may warrant further investigation. In this example, the transcript expression profile from replicate 1 does not agree well with the other two replicates in the control condition. Finally, the DTU classification of the isoforms is encoded as colour or shape, highlighting the changes that are considered significant for the defined criteria. Once created, all plots produced by *RATs* remain customisable via standard ggplot2 operations.

**Figure 4.**
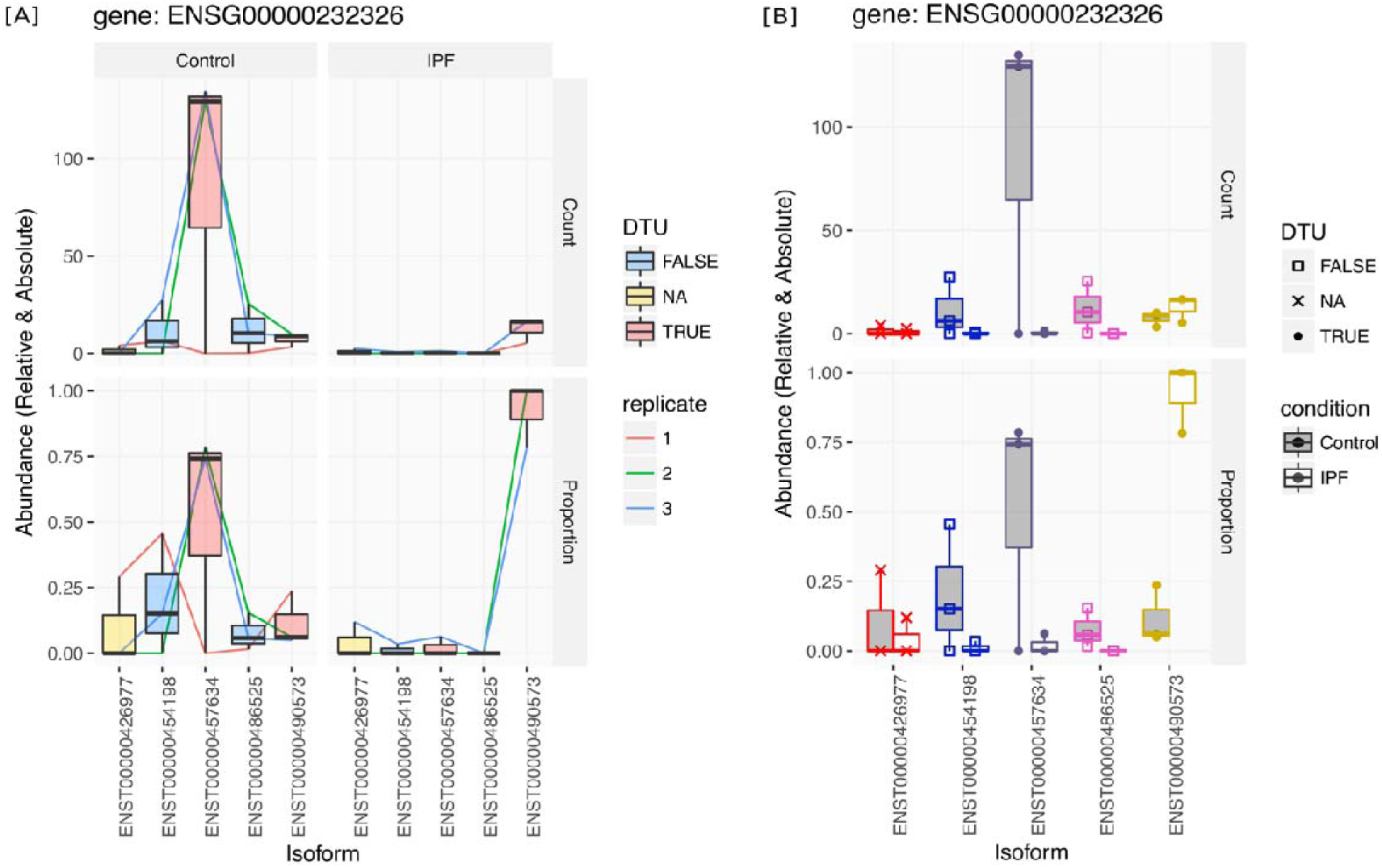
*Isoform abundance plots from RATs for an example gene with five isoforms (Ensembl v60). RNA-seq data was taken from Deng et al.* [28] *as part of the* case study presented in this article. *The isoform expression was quantified with Salmon* [13] *and analysed with RATs. Two conditions are being compared, labelled “Controls” and “IPF” (more details on them in the main text). The upper panels display the absolute abundances (counts) and the lower panels the relative abundances (proportions). The boxplots represent the replicate abundance measurements of each isoform in each condition. In [A], the abundances are organised by condition and colour-filled according to their transcript-level DTU result (blue – non DTU, red – DTU, yellow – not eligible). Coloured lines connect the measurements from each replicate and illustrate the level of consistency among the replicates. In this example, Replicate 1 in the control clearly deviates from the other two. In [B], the abundances are organised and coloured by isoform (red through yellow). The fill-colour of the boxplots indicates the condition (grey – “Controls”, white – “IPF”) while the shape of the points indicates the transcript-level DTU result for the isoform (square – non DTU, cross – not eligible, solid circle -DTU). Style [A] is useful for assessing consistency between replicates and visualising abundance ratio profiles, whereas style [B] facilitates viewing the abundance change of individual isoforms.*

## Identifying DTU in public human RNA-seq data

To test the ability of *RATs* to identify correctly instances of DTU, we compared it against published and validated instances of DTU from publicly available RNA-seq data. We took RNA-seq data from Deng *et al.* [28], who identified non-DGE changes in the isoform levels of genes. The dataset contains 25 million 54-base long single-end Illumina reads per lung tissue sample from three IPF patients and three lung cancer patients as controls. After pre-filtering, Deng *et al.* tested 3098 genes for DTU by quantifying their isoform proportions with RAEM [29] and using Pearsons Chi-squared test of independence with a FDR threshold of 5%. They identified 248 genes that were not differentially expressed but displayed significant DTU. Subsequently, they validated three of them with qRT-PCR: TOM1L1 (ENSG00000141198), CMTM4 (ENSG00000183723), and PEX11B (ENSG00000131779).

As in the original study, Ensembl v60 [30] was used as the source of the reference human genome and its annotation, in which each of the three discussed genes features two isoforms. Unlike the original study, we used Salmon (v0.7.1, with sequence bias correction enabled, 100 bootstrap iterations and default values for the remaining parameters, using *k*=21 for the index) to quantify the isoform abundances. DTU was identified by *RATs* v0.4.4 (with the default parameters *p_thresh*=0.05, *abund_thresh*=5, *dprop_thresh*=0.2, *qrep_thresh*=0.95, and 1000 bootstrap iterations).

With these data and parameters, *RATs* predicted 609 DTU genes according to the gene-level test or 549 DTU genes according to the transcript-level test. 514 DTU genes are common to both methods. The results of *RATs* for the three genes of interest are summarised in Table 1. All three genes exhibited statistically significant changes in isoform proportions, as evidenced by the low p-values for the p, in agreement with the original findings of Deng *et al*. However, only CMTM4 is reported as DTU by *RATs*. This is due to the insufficient reproducibility of the DTU results for TOM1L1 and PEX11B during the DTU bootstrapping process. Of the remaining 245 DTU genes reported by Deng *et al.*, 13 appear in *RATs*’ predictions. The majority of the rejected DTU instances show changes in proportion smaller than our threshold of 0.2, despite high statistical significances, and a also fail to meet our bootstrap reproducibility criterion.

**Table 1.** *Differential Transcript Usage (DTU) results by RATs v0.4.2 for the six transcripts belonging to the three genes identified as DTU by Deng et al. 2013* [28]*. Ensembl v60 was used as the annotation and assembly reference. All transcripts met the effect size (difference in proportion,* ≥*0.2) and significance (p<0.05) requirements and, thus, could be considered DTU. However, only the isoforms of CMTM4 meet the reproducibility threshold (DTU frequency* ≥*95%) and are, therefore, the only ones confidently considered to be DTU by RATs (shaded cells). The corresponding isoform abundance plots by RATs can be found in Supplementary file 2.*

| Gene | Transcript ID | Δproportion | FDR | Bootstrap DTU freq. |
| --- | --- | --- | --- | --- |
| | | ($\times 10^{-2}$) | | |
| TOM1L1 | ENST00000348161 | 25.6 | $1.5 \times 10^{-18}$ | 0.74 |
| | ENST00000445275 | -25.6 | $1.5 \times 10^{-18}$ | 0.74 |
| CMTM4 | ENST00000330687 | 32.1 | $2.5 \times 10^{-130}$ | 1.00 |
| | ENST00000394106 | -32.1 | $2.5 \times 10^{-130}$ | 1.00 |
| PEX11B | ENST00000369306 | -21.0 | $3.4 \times 10^{-34}$ | 0.66 |
| | ENST00000428634 | 21.0 | $3.4 \times 10^{-34}$ | 0.66 |

Repeating the quantification and DTU detection using the current version of the human genome assembly and annotation, Ensembl v87, reveals a different picture of DTU in these data. Despite a 25% increase in the number of annotated transcripts in Ensembl v87 compared to v60 (Table 2), fewer genes were identified as DTU by *RATs*. In total, 511 DTU genes were identified by the gene-level test and 464 by the transcript-level test, with 427 DTU genes identified by both methods. Only 141 DTU genes are in common between our v60 and v87 results (predicted by both test methods), and only 11 of the 248 DTU genes reported by Deng *et al.* are also predicted by *RATs*, the others falling again below our effect size threshold. Additionally, 10,253 gene IDs disappear from v60 to v87 and 15,839 new ones are added. The considerable change of the annotation between the two Ensembl versions affects the number of isoforms the quantification tool can choose from per gene, as well as the multiple testing correction penalty.

**Table 2.** *Expansion of the human annotation between Ensembl v60 and v87. In total, the annotation has 25% more transcript models. The three genes identified by Deng et al.* [28] *(TOM1L1, CMTM4 and PEX11B) have all acquired additional isoform models.*

| Human Annotation | Number of isoforms |  |  |  |
| --- | --- | --- | --- | --- |
|  | Total | TOM1L1 | CMTM4 | PEX11B |
| Ensembl v60 transcripts | 157,480 | 2 | 2 | 2 |
| Ensembl v87 transcripts | 198,002 | 23 | 5 | 3 |

In light of the expanded annotation, the DTU results of the three genes of interest based on the same RNA-seq data changed considerably (Table 3). TOM1L1 no longer demonstrates a switch in primary isoform from ENST00000445275 in the controls to ENST00000348161 in the IPF samples; in fact, none of its isoforms show any significant change in proportion. Furthermore, the isoform abundance estimates have also changed. With Ensembl v60, isoform ENST00000445275 was the dominant isoform in the control samples (Supplementary file 2), whereas with Ensembl v87 it has virtually no expression in either condition. In CMTM4, an abundance shift is still evident, but it occurs between isoforms ENST00000330687 and ENST00000581487 instead of between ENST00000330687 and ENST00000394106. In fact, ENST00000394106 is scarcely detected with Ensembl v87 (Supplementary file 2). Finally, PEX11B demonstrates the same significant change in isoform proportions as it did with Ensembl v60, although this time one of its isoforms (ENST00000369306) does meet the reproducibility criterion and is classified as DTU.

**Table 3.** *Differential Transcript Usage (DTU) results by RATs v0.4.2 for the six transcripts belonging to the three genes identified as DTU by Deng et al.* [28]*. Ensembl v87 was used as the annotation and assembly reference. In bold font are marked the six transcript IDs that were also present in Ensembl v60. The criteria for DTU were a change in proportion* ≥*0.2, p<0.05 and reproducibility* ≥*95%. Transcripts IDs meeting these criteria are shaded light grey. Dark grey shaded cells signify that the isoform’s abundance was below the noise threshold and was not tested. The corresponding isoform abundance plots can be found in Supplementary file 2.*

| Gene | Transcript ID | Δproportion<br>($\times 10^{-2}$) | FDR | Bootstrap DTU<br>freq. |
| --- | --- | --- | --- | --- |
| TOM1L1 | **ENST00000348161** | -0.8 | 0.79 | 0.03 |
|  | **ENST00000445275** | 0.3 |  |  |
|  | ENST00000536554 | -3.0 | 0.14 | 0.03 |
|  | ENST00000536554 | 0 |  |  |
|  | ENST00000570499 | 0.6 |  |  |
|  | ENST00000570623 | 1.3 |  |  |
|  | ENST00000570965 | -1.3 |  |  |
| | ENST00000570965 | -2.6 | $1.5 \times 10^{-3}$ | 0 |
| | ENST00000571319 | 5.6 | $3.8 \times 10^{-4}$ | 0.03 |
| | ENST00000572158 | -7.3 | $1.0 \times 10^{-8}$ | 0.03 |
|  | ENST00000572298 | 0.1 |  |  |
|  | ENST00000572360 | -0.7 |  |  |
|  | ENST00000572405 | -0.1 |  |  |
|  | ENST00000572576 | 0.1 | 0.96 | 0 |
|  | ENST00000572905 | -0.7 |  |  |
|  | ENST00000573607 | 0 |  |  |
|  | ENST00000574318 | 3.1 | 0.01 | 0 |
|  | ENST00000574653 | -3.9 | 0.02 | 0 |
|  | ENST00000574744 | 0 |  |  |
| | ENST00000575333 | 5.6 | $1.3 \times 10^{-3}$ | 0 |
|  | ENST00000575882 | 2.4 |  |  |
|  | ENST00000575909 | 0 |  |  |
|  | ENST00000576932 | 1.3 | 0.21 | 0 |
| CMTM4 | **ENST00000330687** | 28.4 | $2.0 \times 10^{-105}$ | 0.99 |
| | **ENST00000394106** | -1.4 | $2.8 \times 10^{-3}$ | 0 |
| | ENST00000561680 | -1.8 | $2.0 \times 10^{-10}$ | 0 |
| | ENST00000563952 | -1.8 | $1.4 \times 10^{-4}$ | 0 |
| | ENST00000581487 | -23.4 | $6.1 \times 10^{-62}$ | 0.90 |
| PEX11B | **ENST00000369306** | -25.6 | $2.2 \times 10^{-46}$ | 0.96 |
| | **ENST00000428634** | 21.1 | $2.3 \times 10^{-34}$ | 0.71 |
| | ENST00000537888 | 4.4 | $1.3 \times 10^{-08}$ | 0 |

The updates in the genome assembly and annotation are also likely to impact the result of the qRT-PCR validation of the DTU identified in these genes, since PCR primers are designed to target specific transcripts all in a defined annotation. To examine this, the reported primer sequences, designed according to Ensembl v60, were searched for in the respective Ensembl v87 transcript sequences (Figure 5). Two pairs of primers were originally used for each gene [28], each pair consisting of a forward and a reverse primer: One pair was common between the two annotated isoforms of each gene and one was unique to one of them. The unique pair used for TOM1L1 targeted isoform ENST00000445275 in the Ensembl v60 annotation. In the Ensembl v87 annotation, however, these primers match two other isoforms (ENST00000570371 and ENST00000575882). The originally targeted ENST00000445275 has been re-annotated with a truncated 5’ end in Ensembl v87 and now matches only the reverse primer. Thus, it would not be expected to be detected by the given qRT-PCR design. The reverse primer matches an additional isoform in Ensembl v87 (ENST00000570965) that is also annotated with a short 5’ end. Given that isoforms are revised and their ends can change between annotation versions, it is uncertain how many more isoforms may match the primer pair in full or in part. Additionally, the common primers used by *Deng et al.* [28] to detect all the isoforms of the gene match fewer than half of the annotated isoforms in Ensembl v87, and the isoforms captured by the unique primers are not a subset of those captured by the common primers. Interpreting the given qRT-PCR results in the context of the most recent annotation is thus extremely difficult, if not impossible, and any conclusions reached would certainly differ substantially from those published in the original study. For CMTM4, only one pair of primers is listed by *Deng et al.* and it matches two isoforms in Ensembl v87, one of which is the originally targeted ENST00000330687. Finally, in PEX11B, the unique primer pair still uniquely matches isoform ENST00000369306 and the common primer pair matches the new isoform as well as the old ones, lending confidence to the original interpretation of the qRT-PCR quantifications for this gene.

**Figure 5.**
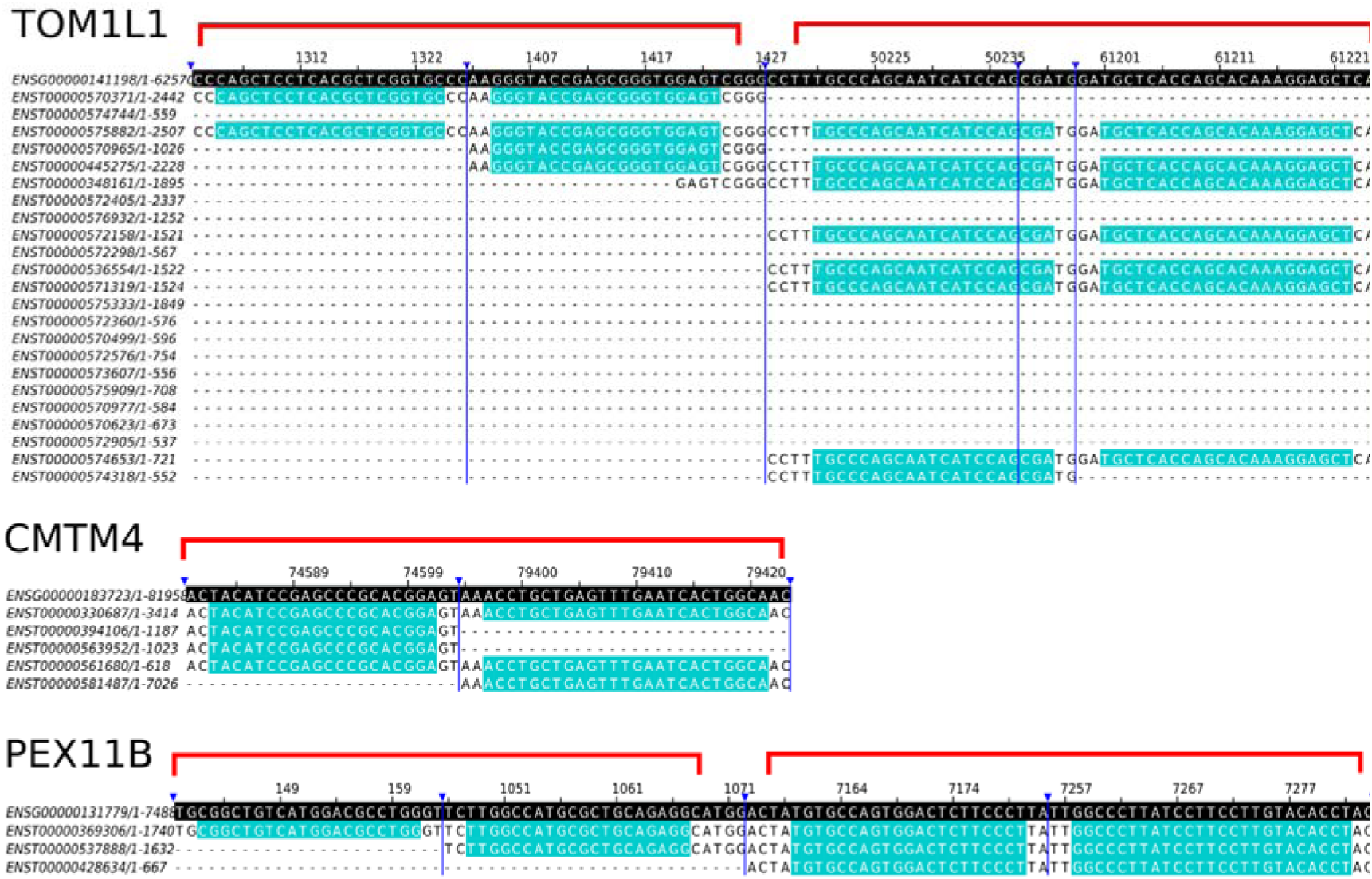
*Sequence matches in Ensembl v87 for the qRT-PCR primers used for DTU validation by Deng et al.* [28]*. The primers were designed based on Ensembl v60. Despite being well designed within the context of Ensembl v60, it is clear in the new annotation that the primers target more transcripts than originally expected. The red brackets indicate the pairs of forward and reverse primers. The dark blue vertical lines indicate portions of the sequence that have been hidden for convenient viewing of the relevant segments. The sequence in black background is the genomic sequence. The primer matches in the isoforms are highlighted in light blue. The alignments were visualized with Jalview* [34].

To verify whether the DTU genes identified by *RATs* were compatible with the known pathology of IPF, we carried out a GO term [31] enrichment analysis for biological processes against the set of all human genes (GO database released 2017-04-24). We used p<0.05, with Bonferroni correction for multiple testing [32]. For the 514 Ensembl v60 DTU genes, we obtained 11 enriched terms (Supplementary file 3) pertaining to regulation of response to stimulus (GO:0048583), regulation of immune system processes (GO:0002682), and symbiosis/parasitism (GO:0044403). Although the exact causes of IPF both in the samples and in general are unknown, viral infections, pollutants and inflammation have been linked to the disease [33]. Basing the GO term enrichment analysis on our 427 Ensembl v87 DTU genes, instead, finds just five enriched terms, pertaining to localisation (GO:0051234, GO:0051179) (Supplementary file 3). No terms were shared between the two enrichment sets, except for the biological process ontology root (GO:0008150). Finally, analysing GO term enrichment for the subset of 141 genes identified as DTU by both annotations resulted in no enriched terms at all.

An alternative resource with which to add context to our results is the Reactome pathway database (Release 59, browser version 3.2) [35–37]. An overrepresentation analysis for the 141 genes identified as DTU by both annotations returned 426 pathway entities spread over multiple clusters. 26 of these entities received p-values <0.05, belonging to the pathway clusters of signal transduction, haemostasis and developmental biology. These entities revolved mainly around the ERBB2 and ERBB4 signalling pathways, while one entity also implicated NOTCH4 signalling. Entities referencing platelet and neutrophil degranulation reinforce the GO terms that implicated an immune response and may relate to foreign objects like viruses or pollutants. The full list of pathways obtained is available in Supplementary file 4.

## Discussion

*RATs* fills a gap in the line-up of differential expression tools, enabling the identification of DTU from isoform expression quantification data. It is based on established statistical methods for ratio comparisons and provides results in formats suitable for both downstream computational analysis and visual inspection. The package includes plotting and summarization routines designed to encode several layers of information, allowing users to examine quickly and easily both the high level DTU picture of their data and to drill down into the individual details and provenance of each result.

The capability of *RATs* to reliably identify DTU depends critically on the upstream isoform expression quantification tools and the quality of the input data they use. This limitation is common to all tools that use an annotation, and any downstream analysis can be affected by the choice of annotation [38]. We showed this by applying *RATs* to publicly available RNA-seq data with validated instances of DTU, using two annotation versions separated by six years. All three validated genes in the study contained additional isoforms in the newer annotation and only one of them displayed similar isoform abundance shifts with both annotations. A second gene showed DTU attributable to different isoforms, depending on the annotation version, whereas the third gene showed no significant DTU with the newer annotation. We also demonstrated that qRT-PCR, often considered the gold standard for differential expression validation, is subject to the same limitation, as evidenced by the multiple hits of the previously assumed unique primer sequences in the newer annotation. qRT-PCR primers are designed to target unique sequences in the transcriptome. However, the transcriptome remains a work in progress even for the most intensively studied model organisms. The natural extensive sequence overlap between isoforms together with the ongoing discovery of additional isoforms may mean that, for many genes, qRT-PCR is not a suitable method for the validation of transcript abundance changes (DTE and DTU alike).

For hybridisation-based methods like qRT-PCR to serve as a reliable validation method of RNA quantification, the suitability of the primers should first be validated by sequencing the captured transcripts. Additionally, it has been reported that pre-filtered annotations improve quantification performance [39], and are likely to be helpful in primer design as well. Such annotations could be obtained by including a parallel set of full-length isoform RNA-seq data in the experimental design, such as via PacBio sequencing (http://www.pacb.com) or Oxford Nanopore Direct RNA-seq (https://nanoporetech.com). An additional advantage of this approach is that it effectively defines the relevant transcriptome for the specific experiment [40–43]. This may be of importance for experiments focussing on specific tissues or developmental stages of an organism where the transcriptome of the sample is likely to be only a subset of the global reference transcriptome of the organism.

Despite these concerns over the annotation, DTU results by *RATs* were shown to be enriched for GO terms that may be relevant to the symptoms and risk factors of IPF or lung cancer. As the controls in the study were not “healthy” individuals, it is likely that the DTU results from this comparison include genes related to the cancer pathology in addition to those associated with the IPF pathology. It is worth keeping in mind that DTU analysis only considers shifts in the ratios of isoforms and that there may be additional genes and transcripts differentially regulated that are not reported here. In fact, no single analysis type among DGE, DTE and DTU can give a complete picture of differential expression; at least two of these analyses must be carried out.

In conclusion, we offer a computational method for identification and visualisation of differential transcript usage and recommend that caution and scrutiny must be exercised in the interpretation of quantifications, whether they be from RNA-seq or qRT-PCR.

## Acknowledgements

This work was supported by Wellcome Trust Strategic Awards [098439/Z/12/Z and WT097945], and Biotechnology and Biological Sciences Research Council Grants [BB/H002286/1; BB/J00247X/1; BB/M010066/1; BB/M004155/1].

